# Energy flux couples sulfur isotope fractionation to proteomic and metabolite profiles in *Desulfovibrio vulgaris*

**DOI:** 10.1101/2023.08.27.555018

**Authors:** William D. Leavitt, Jacob Waldbauer, Sofia S. Venceslau, Min Sub Sim, Lichun Zhang, Flavia Jaquelina Boidi, Sydney Plummer, Julia M. Diaz, Inês A.C. Pereira, Alexander S. Bradley

**Author notes:** Current Address: GenIbet Biopharmaceuticals, Oeiras, Portugal (Sofia S. Venceslau). Instituto Nacional de Tecnología Agropecuaria, EEA Rafaela, Rafaela, Santa Fe, Argentina. Conflicts: The authors declare no conflicts of interest.

## Abstract

Microbial sulfate reduction is central to the global carbon cycle and the redox evolution of Earth’s surface. Tracking the activity of sulfate reducing microorganisms over space and time relies on a nuanced understanding of stable sulfur isotope fractionation in the context of the biochemical machinery of the metabolism. Here we link the magnitude of stable sulfur isotopic fractionation to proteomic and metabolite profiles under different cellular energetic regimes. When energy availability is limited, cell specific sulfate respiration rates and net sulfur isotope fractionation inversely co-vary. Beyond net S isotope fractionation values, we also quantified shifts in protein expression, abundances and isotopic composition of intracellular S metabolites, and lipid structures and lipid/water H isotope fractionation values. These coupled approaches reveal which protein abundances shift directly as a function of energy flux, those that vary minimally, and those that may vary independent of energy flux and likely do not contribute to shifts in S-isotope fractionation. By coupling the bulk S-isotope observations with quantitative proteomics, we provide novel constraints for metabolic isotope models. Together, these results lay the foundation for more predictive metabolic fractionation models, alongside interpretations of environmental sulfur and sulfate reducer lipid-H isotope data.

## 1. INTRODUCTION

Sulfate reducing microorganisms (SRMs) play a central role in Earth’s carbon and sulfur cycles. Sulfate respiration by microorganisms has played a role in the Earth’s carbon cycle for billions of years, ever since sulfate became an abundant oxidant circulated by oceans and rivers (Fike *et al*., 2015; Lloyd *et al*., 2020). Extant SRMs degrade up to half the organic carbon in seafloor sediments (Jørgensen, 1982; Bowles *et al*., 2014; Jørgensen *et al*., 2019), and are integral in balancing Earth’s oxidant budget through the production of iron sulfides (Berner and Canfield, 1989; Hayes and Waldbauer, 2006). Most marine sediments where sulfate reducers are active persist under severe energy limitation (Jørgensen and Marshall, 2016; Bradley *et al*., 2020; Røy *et al*.). Tracing the metabolic activity of SRMs across energy gradients in nature is challenging given their rare doubling and low biomass. The distribution of natural abundance sulfur and hydrogen isotopes in the inorganic (sulfate and sulfide) and organic (lipid) byproducts of SRMs provides a record of microbial metabolic activity long after the environment and community have vanished. Moreover, little is known about sulfate reducing bacterial protein dynamics in the face of chronic energy limitation. To address these limitations, we quantified the S isotope, proteome, lipid abundance and lipid H-isotope phenotypes of energy limited sulfate reducers cultivated steady-state.

The majority of microbial sulfate reduction (MSR) occurs in marine sediments where energy supply is limited by the input of organic carbon, driving microbes in the system toward a state of persistent starvation (Goldhaber and Kaplan, 1975; Lever *et al*., 2015; Jørgensen *et al*., 2019; Bradley *et al*., 2020; LaRowe *et al*., 2020). Most laboratory MSR experiments, by contrast, are performed in closed (‘batch’) systems under conditions of energy and nutrient excess, such that most experiments for S isotope or proteomic analysis are biased toward energy and nutrient replete and closed-system conditions. We do not know yet what occurs within the metabolic network of SRMs under the range of energetic regimes in either the lab or nature (Sim *et al*., 2023). This hinders the interpretive value of stable isotopes and metaproteomes in ecosystems where SRMs are major players.

Energy availability is a key determinant of microbial activity and the vigor with which biogeochemical cycles operate. Empirically derived energy flux versus isotope fractionation relationships (Sim *et al*., 2011; Leavitt *et al*., 2013) establish the basis for a range of models that incorporate sulfate reducer biochemistry and in vitro observations to sulfate reducer fractionation models (Bradley *et al*., 2011, 2016; Wing and Halevy, 2014; Wenk *et al*., 2017), yet many of the core assumptions in such models remain untested. In this study, we quantified shifts in the proteome of a model sulfate reducer under different energy availability, where sulfur isotope fractionation co-varied with energy flux and cell specific sulfate reduction rate (csSRR; Figure 1). Given the central role of empirical data in building and testing such models, we sought to track the biochemical (proteomic and electron carrier) responses to differences in energy flux. In particular, under different electron-donor limited steady-state growth rates, we ask: Do central metabolism protein abundances increase, decrease, or remain constant? Do intercellular reservoirs of sulfate reduction intermediates respond to change? Does the abundance or identity of electron carriers change at high versus low energy flux? In addressing these questions, we provide a set of new constraints for the next generation of isotope-metabolism models, which in turn will enable a more robust interpretation of sulfur isotopic records across environments and time periods.

**Figure 1.**
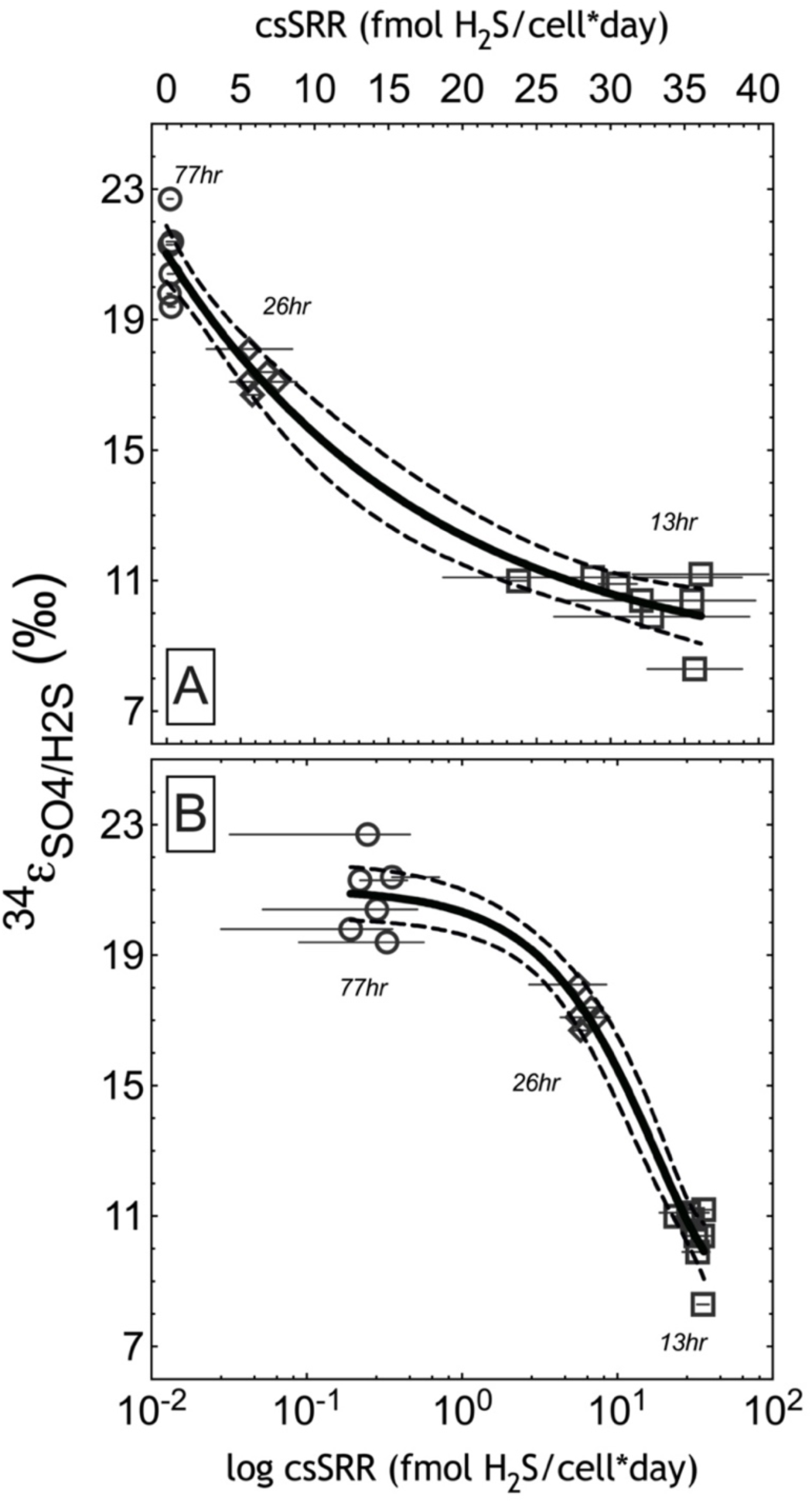
Sulfur isotope fractionation under three csSRR under electron-donor limitation. Either the (A) linear or (B) logarithmic x-axes of csSRR versus net isotope fractionation between residual sulfate in the effluent and accumulated product sulfide (y-axis). The nonlinear regression (Equation 3) was applied following Leavitt et al. 2013, in order to calculate the empirical fractionation limit (^34^ε_sulfate/sulfide_) (r^2^ = 0.97) as a function of csSRR. The best-fit estimate (solid line) with 95% confidence intervals (dashed lines). The Y-intercept is 21 ± 0.8 ‰, Y-plateau is 8.3 ± 1.3 ‰, and the *k*_ε_ = 0.056 ± 0.015. The 77hr and 13hr endmember experiments are new to this study, while the 26hr data recalculated from Leavitt et al. 2019.

## 2.0 METHODS

The model sulfate reducing bacterium *Desulfovibrio vulgaris* DSM644^T^ (WT) was cultivated following established protocols (Leavitt, Venceslau, *et al*., 2016), with the following modifications: chemostat media contained 30 mM lactate as the electron donor and 30 mM sulfate as the electron acceptor, at 30.5 ± 0.1°C and pH 7.2 ± 0.1. Axenic populations of DSM644 grew and reduced sulfate at three distinct electron donor fluxes, where sulfate and other nutrients (P, N, trace metals, vitamins) were provided in excess and lactate influx limited growth and respiration. Samples were taken after bioreactors completed at least five turnovers, and had achieved steady-state. The bioreactor (chemostat) systems are identical to those utilized recently for other DvH experiments (Leavitt *et al*., 2019). The slowest (77hr turnover time) and fastest (13hr turnover time) experiments were performed in duplicate parallel reactors for each rate, with a number of intervals sampled for bulk S isotopes, while for proteomics and cellular metabolites, samples are drawn only at one interval for each reactor (in technical replicate). For the intermediate rate (26hr turnover time), the experiments were performed in a single reactor, sampled at different turnover internals for both isotopes and proteomics, as previously reported (Leavitt *et al*., 2019). The sampling regime, analytical measurements and calculations followed those in other recent sulfate reducer chemostat studies (Leavitt *et al*., 2013, 2019; Bradley *et al*., 2016), summarized below.

### 2.1 Chemostat calculations

Dilution rate (*D*) of the chemostat reactor volume is calculated as:

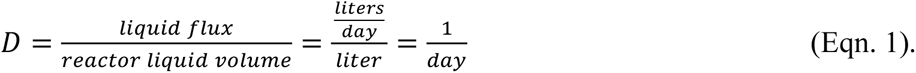

The csSRR in an open system, where sulfate flux values may be substituted into Equation 2, when flux balance is conserved:

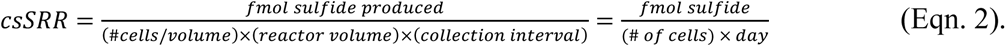

### 2.2 Major reactant and product analyses

Extracellular sulfate and lactate samples were stored at –20 °C until analysis. Lactate concentrations in fresh and spent media were determined by an Agilent 1100 HPLC (Agilent Technologies, Santa Clara, CA, USA) on an Aminex 87H column (Bio-Rad, Hercules, CA, USA). The sample was eluted with 8 mM sulfuric acid as an isocratic mobile phase at a flow rate of 0.6 ml/min, and the column effluent was passed through UV-visible and refractive index detectors. Sulfate concentrations in fresh and spent media were determined on a Dionex DX 500 ion chromatograph (IC) equipped with an AS11-HC column (Dionex, Sunnyvale, CA, USA), using a gradient elution with KOH as mobile phase. After electrochemical suppression, quantification was achieved with a conductivity detector.

For sulfide quantifications, the zinc acetate preserved samples (see trapping procedure in Leavitt et al. 2013) from each sample was quantified using a modified Cline method, as done previously (Cline, 1969; Leavitt *et al*., 2013). Analytical grade sodium sulfide (>98.9% Na2S*9H2O) was used as the standard, and prepared in deoxygenated (boiled and N2-sparged) growth medium, by precipitating the sodium sulfide with excess zinc acetate (anhydrous), mimicking the detailed protocol already published (Leavitt *et al*., 2014), with chemostat zinc-trap procedures following other recent chemostat studies (Leavitt *et al*., 2019).

### 2.3 Bulk sulfur isotope analyses and S isotope notation

Bulk sulfur isotope analyses were performed as on the medium sulfate, reactors sulfate and trapped sulfide following a recent study (Leavitt *et al*., 2019). Samples for bulk S isotopes were the medium sulfate before it was allowed to enter the reactors, residual dissolved sulfate removed from the reactors and after the reactors, and the microbialgenic sulfide, removed from gas traps. The fractionation between sulfate and sulfide is calculated as ^34^ε from the measured δ^34^S values measured on trapped sulfate and sulfide. The variability in ^34^S/^32^S of a distinct S-bearing species or operationally defined pool, *y*, is reported as δ^34^Sy (Equation 3a). The difference between two pools, such as sulfate (A) and sulfide (B) is Equation 3b (*y* = A or B), and are reported in alpha (Eqn. 4a) or epsilon (Eqn 4b) notation, where the latter is in permille.

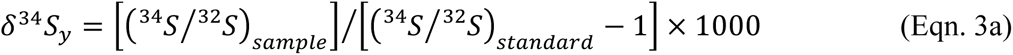

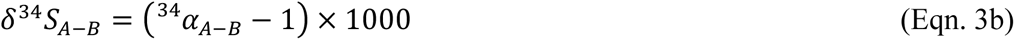

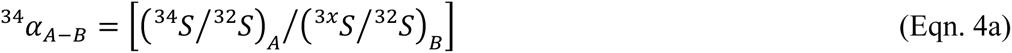

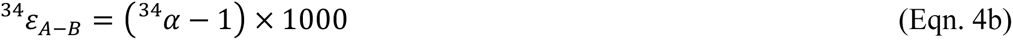

### 2.4 Rate--fractionation model

We applied the non-linear regression one-phase decay model to the net S-isotope fractionation and specific rate estimates developed in a prior study (Leavitt *et al*., 2013). Results of the model fit and standard error estimates, fitting parameters and their standard errors, and the 95% confidence intervals (dashed lines) are presented in Figure 1. The fitted rate constant is first order with respect to the rate limiting reactant (electron donor, lactate). This assumption is valid because the growth and respiration of *D. vulgaris* chemostat populations are limited by electron donor influx, whereas terminal electron acceptor sulfate is always present in excess. The non-linear rate/fractionation model is presented in equations 5 and 6.

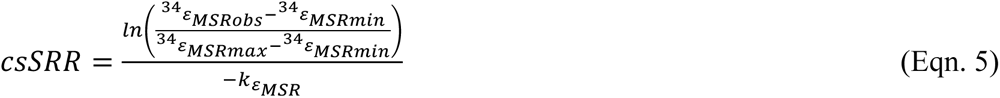

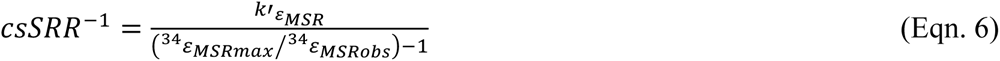

Data inputs are the measured (observed) ^34^ε*_MSRobs_* or for a given dilution rate (*D*) and the corresponding csSRR. Fitting parameters are as follows: *k*, a pseudo-first order rate constant specific to each non-linear regression (*k*ε, *k*’ε,); ^34^ε*_MSRmax_*, the theoretical maximum fractionation as *csSRR^-^*^1^ approaches infinity (which physically corresponds to *D* approaching zero), and in the case of the one-phase decay model (Eqns. 3, 4) the ^34^ε*_MSRmin_* corresponds to classically defined ‘plateau’, where the minimum value corresponds to the rate-limiting step in sulfate reduction when *D. vulgaris* grows and metabolizes at its *µ*max.

### 2.5 Intracellular S-metabolite abundance and S-isotope analyses

Intracellular sulfur metabolite quantification and S-isotope analysis were performed using preparative ion chromatography coupled to multi-collector inductively coupled plasma mass spectrometer (MC-ICP-MS) at the California Institute of Technology. The experimental and analytical procedures are described in greater detail elsewhere (Sim *et al*., 2017) and summarized here. Biomass samples from the 13hr and 77hr reactors were washed with anaerobic cold saline solution to remove culture medium constituents and flash frozen in liquid nitrogen until extraction. The frozen pellet was thawed and resuspended in 0.6 mL degassed deionized water (DW) under an anaerobic atmosphere of nitrogen:hydrogen (90:10) (Coy Manufacturing Co., Ann Arbor, MI, USA). After mixing with an equal volume of cold methanol, the sample was left on dry ice for 30 min and then thawed on ice for 10 min in 20 µL of 50 mM zinc chloride and 100 µL of 13 mM formaldehyde were added to prevent the oxidation of sulfide and sulfite, respectively. The supernatant was recovered by centrifugation. Sulfur metabolites, sulfate, sulfite, and APS, were quantified by IC using a gradient dilution with KOH as mobile phase, and the eluent fractions corresponding to sulfate and APS were collected for isotopic analysis. Collected APS was quantitatively converted to sulfate *via* hydrolysis under acidic conditions. Then samples containing dissolved sulfate were dried on a hot plate and diluted in 5% nitric acid to a sulfate concentration of 20 µM to match the in-house working standard. S isotope analysis was carried out following the original method (Paris *et al*., 2013) via MC-ICP-MS (Thermo-Fisher Scientific Neptune Plus, Bremen, Germany). Sulfur isotope ratios of the sample and working standard were measured alternatively, and instrumental blank was estimated after each block. The mean blank value was subtracted from the measured signal for ^32^S and ^34^S, and the measured ^34^S/^32^S ratios were calibrated using a linear interpolation between the two bracketing standard values.

### 2.6 Cell associated (in)soluble redox intermediates

Menaquinone and NAD(P)(H) levels in 13hr and 77hr biomass samples were measured at the Proteomics & Mass Spectrometry Facility at the Danforth Plant Science Center, following established methods (Lunn *et al*., 2006; Arrivault *et al*., 2009). Biomass pellets were prepared as noted above for intracellular S metabolite analyses by cold centrifugation and flash-freezing. The nicotinamide cofactors were quantified using a custom LC-MS, an Eksigent MicroLC and Thermo Q-Exactive using a 0.5 x 50 mm Sepax Proteomix SAX column. Samples were extracted using a protocol modified from (Lunn *et al*., 2006). The samples were dissolved in 100 µL water then one microliter was injected onto the LC-MS. The solvents were 25% methanol (B) and 250 mM (NH4)2CO3. The gradient started at 100% B after a hold for 3 minutes the composition was adjusted to 50% over 5 minutes followed by a ramp to 0% B over 3 minutes with a hold for one minute then back to 100% B over 2 minutes followed by a re-equilibration for 12 minutes; the flow rate was 15 µL per minute. Data were recorded in negative profile mode from m/z 200-1000 with a resolution setting of 70,000 (at m/z 200). Peak areas were extracted and normalized against the area for MOPS (internal standard).

A total of six samples were analyzed for the presence of menaquinones MK-5 and MK-6 using MK-4 as a standard. These compounds were analyzed from the organic fraction of the extraction above using a custom Eksigent MicroLC and Thermo Q-Exactive using a 0.5 x 100 mm custom packed PLRPS column. The solvents were water with 0.1% formic acid (A) and acetonitrile with 0.1% formic acid (B). The gradient started at 30% B after a hold for three minutes the composition was adjusted to 100% B over three minutes with a hold at 100% B for two minutes followed by a ramp back to 300% B over two minutes followed by a re equilibration for six minutes; the flow rate was 15 µL min^-1^. Data were recorded in positive profile mode from m/z 400 to 700 with a resolution setting of 140,000 (at m/z 200). These quantities are measured relative to the internal analytical standard (MK-4), then normalized for cell numbers, such that sample size or cell number is not the explanatory variable.

### 2.7 Cell counting

Cell counts were conducted on a Guava EasyCyte HT flow cytometer (Millipore). Flow cytometry samples were preserved in 0.5% glutaraldehyde (final concentration) and stored at –80℃ prior to analysis. For analysis, samples were diluted (1:100) with filtered seawater (0.01 μm),stained with SYBR Green I (Invitrogen) according to the manufacturer’s instructions, and incubated in a 96-well plate in the dark at room temperature for 1 hr. Samples were analyzed at a low flow rate (0.24 μL s^-1^) for 3 min. Cells were gated and counted based on diagnostic forward scatter versus green fluorescence signals. Instrument-specific beads were used to validate performance of the cytometer prior to sample analysis.

### 2.8 Proteomics

Samples for proteomics were collected and analyzed as in (Leavitt *et al*., 2019), with quantitation by diDO-IPTL *in vitro* peptide isotope labeling (Waldbauer *et al*., 2017). The false discovery rate (FDR) for peptide-spectrum matches was controlled by target-decoy searching to <0.5%, using the translated genome of *Desulfovibrio vulgaris* DSM 644^T^ obtained from MicrobesOnline (http://www.microbesonline.org/) and Integrated Microbial Genomes (IMG; http://img.jgi.doe.gov/) as a search database. Protein-level relative abundances and standard errors were calculated in R using the Arm post processing scripts for diDO-IPTL data (Waldbauer *et al*., 2017b; github.com/waldbauerlab).

Significantly differential protein expression between experimental conditions was determined using a *Z*-score for protein abundance differences by taking the difference in the mean (log2-transformed) protein abundance between conditions and dividing it by the sum of the total uncertainty estimate for that protein in the two conditions. This total uncertainty estimate for a given condition, was taken as the root-square sum of the standard deviation of a protein’s abundance across the biological replicates of that condition plus the average standard error of the protein’s abundance across quantified spectra within each replicate. These *Z*-scores were converted to *p*-values assuming a standard normal distribution and then the familywise error rate for significantly differential expression between conditions was controlled using the *q*-value method to correct for multiple testing (Benjamini and Hochberg, 1995).

### 2.9 Fatty acid abundances and lipid hydrogen isotope compositions

Cells from each of the 13hr and 77hr reactors were pelleted and freeze dried after sampling by cold centrifugation. Fatty acid extraction, derivatization to fatty acid methyl esters (FAMEs) and analyses performed following other studies on SRMs (Leavitt, Flynn, *et al*., 2016; Leavitt, Venceslau, *et al*., 2016). The FAME retention times and peak areas were determined by gas chromatography (GC) with thermal conductivity detection (TCD), the δ^2^H values of lipids measured by GC high temperature conversion isotope ratio mass spectrometry (GC-HTC-IRMS) following recent studies (Leavitt, Flynn, *et al*., 2016; Leavitt, Venceslau, *et al*., 2016). Each fatty acid from each sample was measured for δ^2^H a minimum of 3x, for a total 3-technical replicates on each of 2x biological replicates per sampling interval, per each reactor per rate. Hydrogen isotope compositions of medium waters were measured following established methods (McFarlin *et al*., 2019; Taenzer *et al*., 2020). The variability in ^2^H/^1^H of a measured pool, *y*, is reported as δ^2^Hy (Eqn. 7) reported relative to V-SMOW (Vienna Standard Mean Ocean Water). Similar to differences in S-isotope pools, the H-isotope fractionation is reported between medium water and a specific lipid or the weighted abundance of all lipids in a given can be reported in alpha (Eqn. 8b) or epsilon (Eqn. 8b) notation.

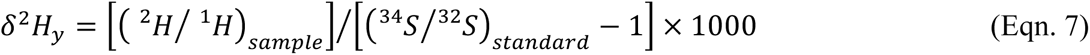

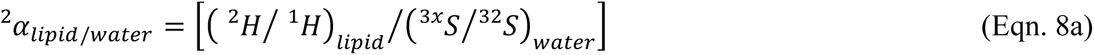

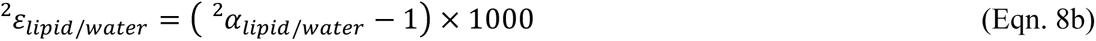

## 3.0 RESULTS

In this study we cultivated model sulfate reducer *Desulfovibrio vulgaris* DSM644^T^ under three different steady-state conditions each limited by electron donor (Figure S1). From all conditions we quantified net sulfate/sulfide S isotope fractionation and protein-level gene expression, while from the fastest (13hr) and slowest (77hr) conditions we also quantified the intracellular sulfate S isotope composition, cellular S metabolite abundances, lipid/water H isotope fractionation, electron carrier abundances,. Each chemostat steady-state and related rates yielded statistically unique responses in each category.

### 3.1 Sulfate reduction rates, S-isotope fractionation, and core S-metabolites

In each experiment sulfate reduction rate scaled with bioreactor turnover time, and in turn, net sulfur isotope fractionation between reactant sulfate and product sulfide scaled with cell specific sulfate reduction rates. The average reactor turnover times (τ) were 13 ±1 hours, 26 ±2 hours, and 77 ±5 hours, which corresponded to estimated cell-specific sulfate reduction rates (csSRR) of 31.9 ±5.6, 6.2 ±1.4, and 0.3 ±0.2 fmol per cell per day (Figure 1). The average sulfur isotope fractionation values between reactant sulfate and product sulfide, ^34^εSO4/H2S, were: 21 ±1 ‰, 17±1‰, and 10 ±1‰, from slowest to fastest, respectively (Figure 1), and are significantly different between the three rates. A non-linear one-phase decay model fit the rate versus fractionation relationship, following the model in Leavitt an colleagues, though the maximum and minimum values are unique to this dataset, relative to those in the literature (Chambers *et al*., 1975; Sim, Bosak, *et al*., 2011; Sim, Ono, *et al*., 2011; Leavitt *et al*., 2013).

The abundance of key sulfate reduction intermediates scales with electron donor flux. The relative abundance of intracellular metabolic intermediates from the fast (τ = 13hr) and slow (τ = 77hr) populations tracked net rate. Specifically, the relative quantities of biomass-associated (intracellular) sulfate, APS and sulfite. The intracellular sulfate, APS, and sulfite levels were consistently higher in the fastest (13hr) relative to slowest (77hr) doubling populations, respectively (Figure 2A). Intracellular sulfate-S isotope compositions were measured at the fastest and slowest turnover times (Figure 2B). Intracellular sulfate S-isotope values are enriched by 6.5 and 7.7 ‰, relative to the residual sulfate in the reactors, from the fast turnover times (τ = 13hr) reactors. In the slowest turnover time experiments (τ = 77hr), intracellular sulfate S isotope values were only 1.1 and 1.2 ‰ enriched relative to residual reactor sulfate (Figure 2B). The directionality of this enrichment is consistent with a recent suite of batch experiments (Sim *et al*., 2017). Rate and model calculations are detailed in the Methods.

**Figure 2.**
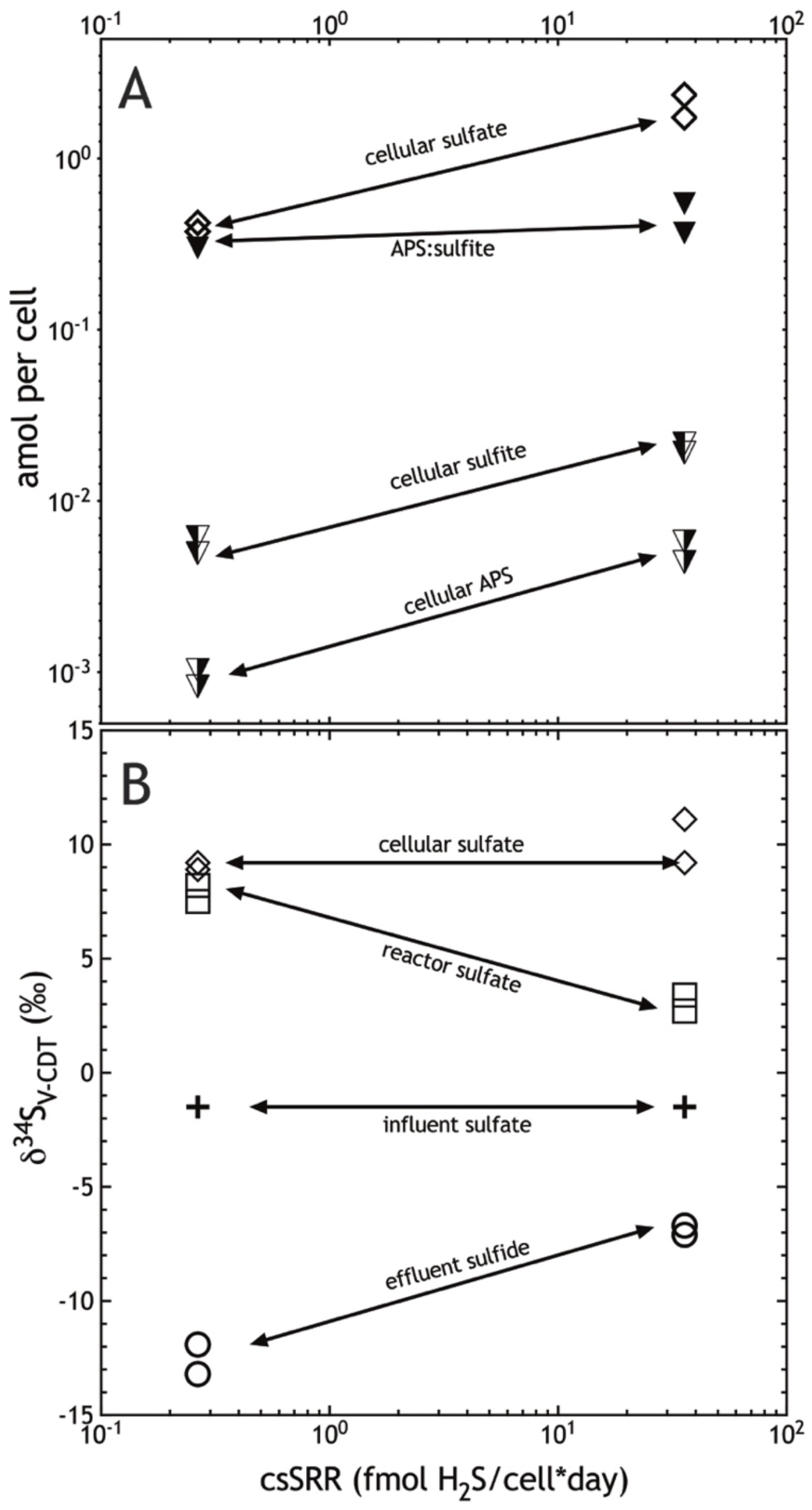
Sulfur isotopes and S metabolites from the slowest (77hr) and fastest (13hr) reactors. (A) attamol per cell quantities of intracellular sulfate, APS, and sulfite, as well as the APS:sulfite per cell ratio. (B) Sulfur isotope compositions of reactant (influent) sulfate, intracellular sulfate, residual reactant (reactor) sulfate, pooled product effluent) sulfide; and the off-set between reactor sulfate and effluent sulfide in (B) gives the estimates of ^34^ε_sulfate/sulfide_ in Figure 1.

### 3.2 Lipid abundance, lipid H-isotope values, lipid and soluble electron carriers

To determine if electron donor limitation is recorded in membrane lipid profiles or their and H-isotopic compositions, we measured the relative abundance of different fatty acids (FA), the hydrogen isotope composition of those FAs, and the menaquinone and NAD(P)(H) abundances from the 13hr and 77hr rate regimes.

Each *D. vulgaris* population produced abundant FAs with differing quantities of *iso-*, *anteiso-* and *n*-fatty acids between 15 to 19 tail carbons, with 0 to 1 double-bonds, and small quantities of 3-hydroxy-C18:0. The faster growing *D. vulgaris* produced FA profiles composed of about 35% *iso*-C17:1, with each other FA making up no more than 10% each, whereas the 77hr membranes were composed of ∼25% *iso*-C17:1, with ∼17% *n*-C18:1 (Figure 3). The largest differences in FA abundances between the 13hr and 77hr *D. vulgaris* populations were shifts in *iso*-C17:1 and in *n*-C18:1 (Figure 3). Meanwhile, the hydrogen isotopic composition of the different fatty acids captured a consistent pattern between the fastest versus the slowest populations. Overall a small, but reproducible shift was apparent, with more depleted values in the faster 13hr cells relative to the slow 77hr cells. The mass-weighted average fractionation between lipids and growth water (^2^ε*l/w*_*total*) for the 77hr cells is –158 ‰, as opposed to –175 ‰ for 13hr cells (2σ = 8 ‰). The compound-specific fractionation values (^2^ε*lipid/water*) for each individual lipid was also greater in the slowest relative to fastest growing cells (Figure 3B). The most depleted lipid for each rate condition was *anteiso-*C17:1, where the most enriched were *n-*C17:1 and *n-*C18:1. The isotopic ordering of individual lipids (^2^ε*lipid/water*) was similar for both populations (Figure 3B).

**Figure 3.**
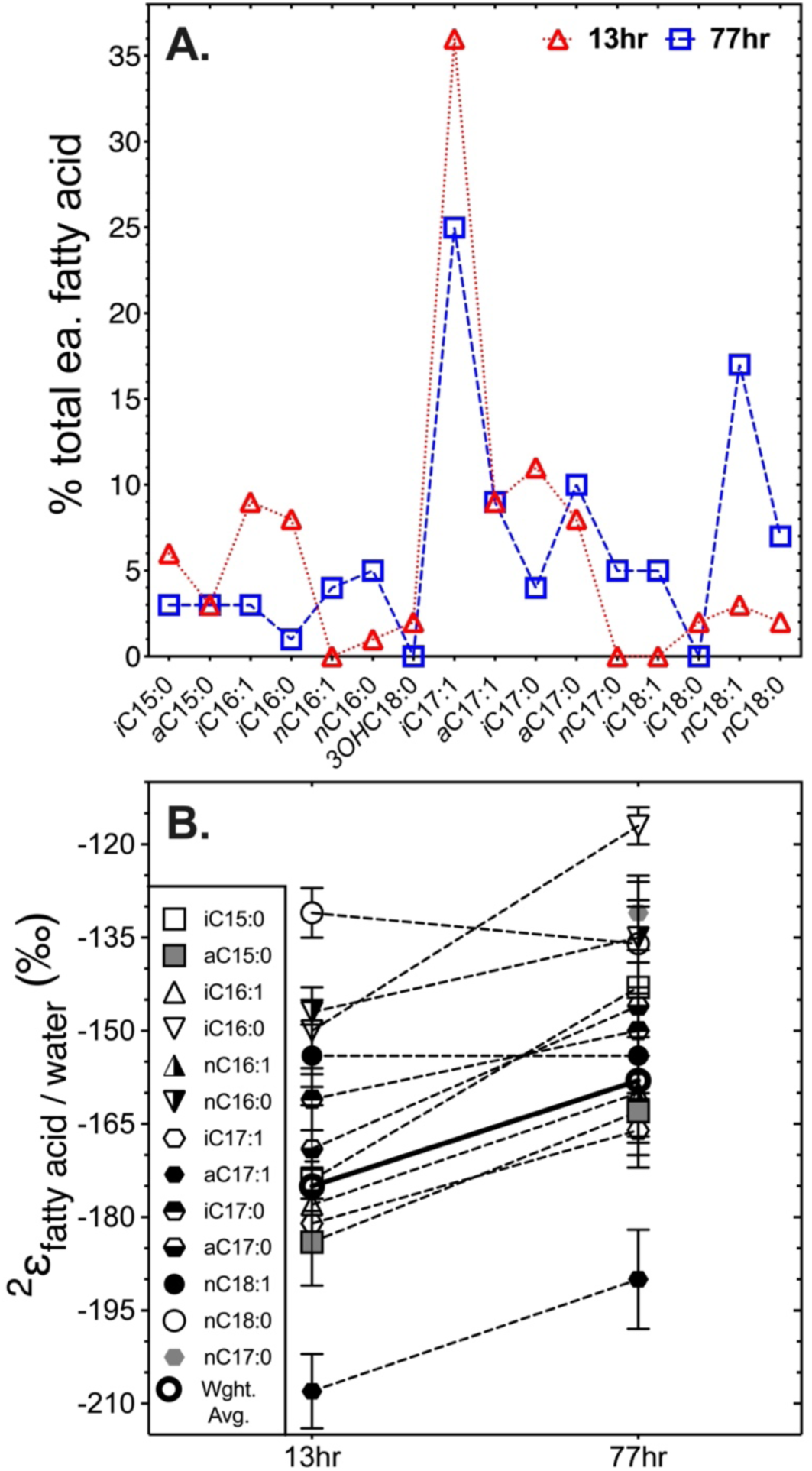
The *Dv*H lipid abundance and lipid/water H-isotopes. (A) Fatty abundances from the 13hr (red dotted line and triangles) and 77hr (blue dashed line and squares). (B) The ^2^ε*_lipid/water_* values for individual fatty acids, as well as the abundance the weighted average (heavy open circle) for the 13hr vs. 77hr chemostat grown cells.

The relative quantities of membrane associated electron carriers menaquinone 5 and 6 (MK-5, – 6), and soluble electron transfer intermediates NADP(H) and NAD(H), were measured on biomass from the τ = 13hr and 77hr reactors. More NADH was found in τ = 13hr relative to 77hr cells, and NADPH was only detectable in 13hr cells (Figure S2A). Similarly, more of both MK’s were present in the 13hr cells (Figure S2B), and while NAD(P)(H) levels were more variable between replicates, likely due to our sampling strategy, the general trend showed more of all oxidized H-shuttles, NAD^+^ and NADP^+^ in the 13hr cells, which were both orders of magnitude more abundant than the reduced counterparts in both populations. Future applications of these measurements require a more robust method to generate high fidelity estimates (García *et al*., 2018).

### 3.3 Proteomic response to changes in energy flux

To identify how protein expression modulates energy metabolism at each electron-donor flux, we quantified the whole proteome response in the *D. vulgaris* population at each steady-state growth rate. We quantified 387 proteins at all rates, in all replicate reactors, across all sampling intervals (Figure S3), representing 11% (387 of 3535) of all predicted protein-coding genes in *Desulfovibrio vulgaris* DSM 644^T^. To understand how protein expression patterns and COGs correlate to the magnitude of sulfur isotope fractionation, we focus on proteins where abundance increased or decreased significantly *with* or *inverse* to csSRR (*q* < 0.05). Of the aforementioned 387 proteins, 37 increased in abundance as csSRR increased (more in 13hr > 26hr > 7 h), versus 28 that increased in abundance inverse to csSRR (most abundant in 77hr > 26hr > 13hr). Of the 37 up with rate, 11 are from ‘energy production and conversion’ COG category, 12 from translation and ribosome production, 7 of unknown function and the other 7 across four other COG categories (see Table 1).The large number of ribosomal proteins in fast (13hr) populations is consistent with prior observations, where the relationship between bacterial cell growth, translation rates, and ribosome abundance covary (Klumpp *et al*., 2013; Bosdriesz *et al*., 2015). From the 28 the scaled inverse to rate, only 2 were from translation and ribosome biogenesis, whereas 9 are from energy production and conversion, four from unknown function, with the remaining 13 scattered across other COG identifiers (Table 1). From those same 65 proteins, a clear secondary COG function was also identified for 5 of these (Table 1). Below we focus primarily on the energy and carbon metabolism proteins (COG category C) that are most likely involved in catabolic S and anabolic H isotope fractionation. These proteins are summarized in Figure 4 (and Figure S4) and discussed below.

**Figure 4.**
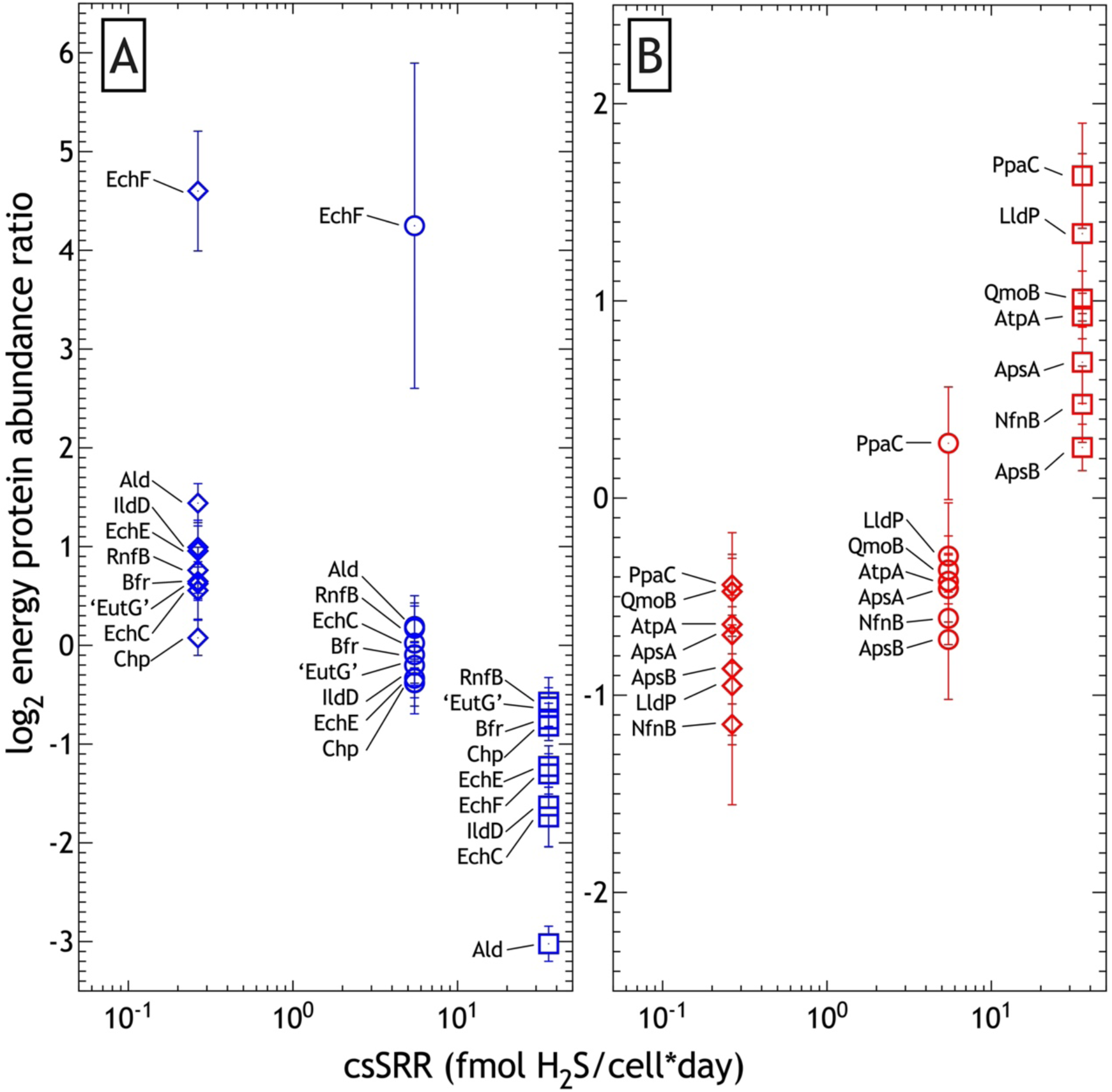
The core sulfate reduction and energy metabolism proteins in *Dv*H that vary with csSRR. (A) Energy proteins that decrease in abundance as reductant-limited growth and csSRR increases, or (B) energy proteins that increase in abundance as growth and csSRR decreases. The y-axes are plotted on the same scale in Figure S4.

**Table 1.**
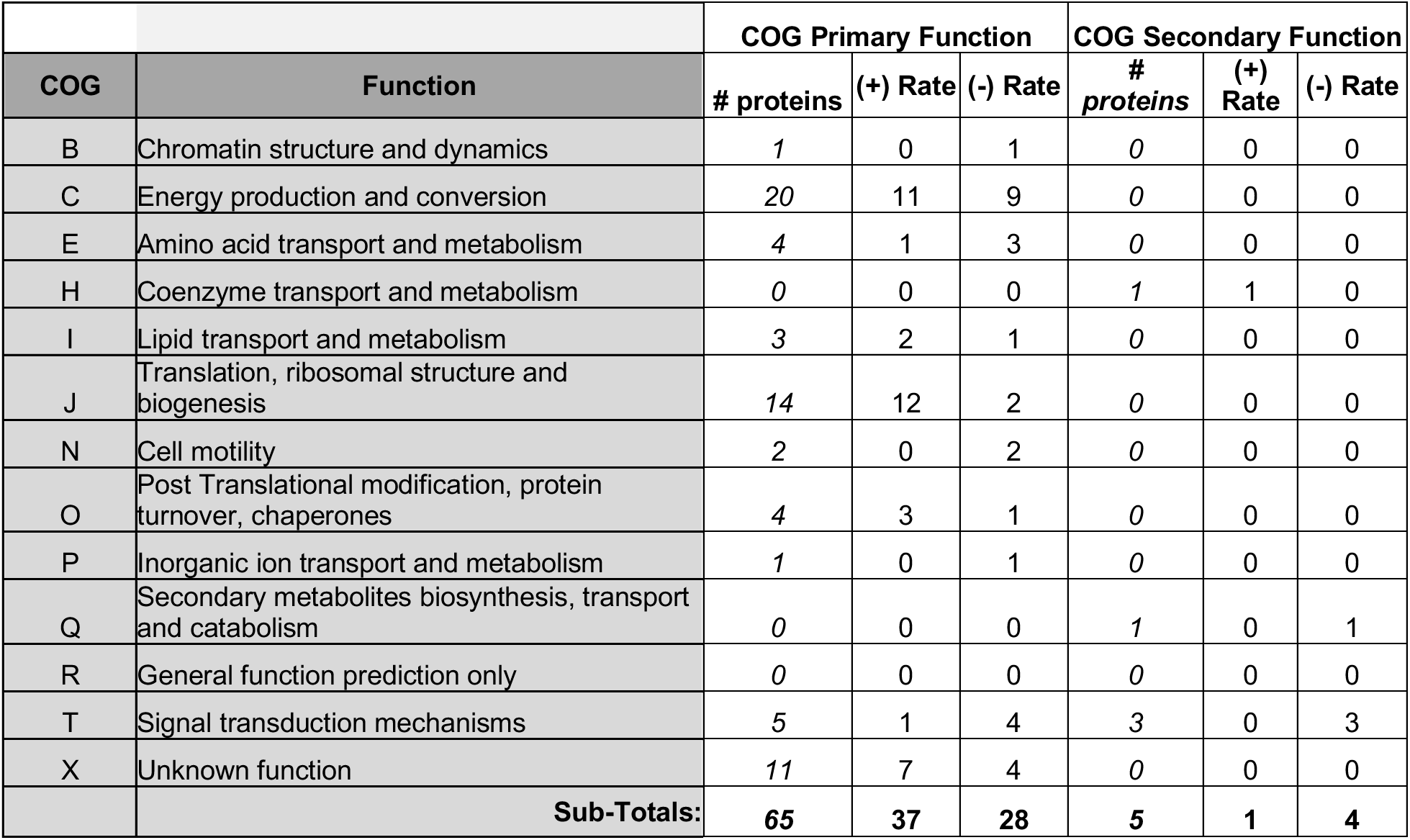
The differentially expressed proteins in *Dv*H with rate, by COG. The number of proteins by COG category increased or decreased consistently with electron donor regulated rate. (+) positive abundance scaling with csSRR vs. (–) inverse abundance with csSRR.

| COG | Function | COG Primary Function |  |  | COG Secondary Function |  |  |
| --- | --- | --- | --- | --- | --- | --- | --- |
|  |  | # proteins | (+) Rate | (-) Rate | # proteins | (+) Rate | (-) Rate |
| B | Chromatin structure and dynamics | 1 | 0 | 1 | 0 | 0 | 0 |
| C | Energy production and conversion | 20 | 11 | 9 | 0 | 0 | 0 |
| E | Amino acid transport and metabolism | 4 | 1 | 3 | 0 | 0 | 0 |
| H | Coenzyme transport and metabolism | 0 | 0 | 0 | 1 | 1 | 0 |
| I | Lipid transport and metabolism | 3 | 2 | 1 | 0 | 0 | 0 |
| J | Translation, ribosomal structure and biogenesis | 14 | 12 | 2 | 0 | 0 | 0 |
| N | Cell motility | 2 | 0 | 2 | 0 | 0 | 0 |
| O | Post Translational modification, protein turnover, chaperones | 4 | 3 | 1 | 0 | 0 | 0 |
| P | Inorganic ion transport and metabolism | 1 | 0 | 1 | 0 | 0 | 0 |
| Q | Secondary metabolites biosynthesis, transport and catabolism | 0 | 0 | 0 | 1 | 0 | 1 |
| R | General function prediction only | 0 | 0 | 0 | 0 | 0 | 0 |
| T | Signal transduction mechanisms | 5 | 1 | 4 | 3 | 0 | 3 |
| X | Unknown function | 11 | 7 | 4 | 0 | 0 | 0 |
| Sub-Totals: |  | 65 | 37 | 28 | 5 | 1 | 4 |

## 4.0 DISCUSSION

In this study we show how environmental conditions influence inorganic (^34^εSO4/H2S) and organic (^2^εlipid/water) biosignatures with the model sulfate reducer, *D. vulgaris*. Experimentally imposed shifts in cell specific growth and metabolic rates (csSRR) due to different electron donor flux yield clear shifts in key protein and metabolite abundances, lipid abundance and H-isotope values, cellular S-metabolites and net S isotope values. While the experiments herein cover a smaller range of csSRR and concomitant ^34^εSO4/H2S than prior chemostat studies (Chambers *et al*., 1975; Sim, Bosak, *et al*., 2011; Leavitt *et al*., 2013), the novel suite of tools we employ to quantitatively track physiological (proteomic, metabolite), organic (lipid/water), and inorganic (sulfate/sulfide) isotope responses provide a new window into how sulfate reducing bacteria operate at different energy availabilities.

### 4.1 Energy flux drives shifts in key proteins and metabolites

The abundance of some energy metabolism proteins is clearly coupled to shifts in csSRR and ^34^εSO4/H2S (Figure S4). Some, but not all, core sulfate respiration and energy metabolism proteins scaled positively or negatively with energy flux (Figure 4). The *proteins that decreased* as *energy flux decreased* (Figure 4B) include those integral to APS reduction to HSO3^-^/SO ^2-^ (ApsAB, QmoAB, PpaC) – together these proteins are part of the Rex regulon, which is a repressor of sulfate adenylyltransferase that is regulated by NADH (Christensen *et al*., 2015).

Given that APS reduction is the first step in MSR to link sulfate respiration to membrane-bound electron transport (and hence energy conservation) via the Qmo complex, this reveals the cellular scale coupling of MSR proteins to environmental limitation in electron donor, and the consequences for intracellular S intermediates and isotopes (Figure 2). On the anabolism side, part of the NADPH/NADH electron-bifurcating transhydrogenase (NfnB) was much less abundant at low csSRR (Figure 4B). Nfn likely plays a significant role in sulfate reducer lipid-H isotope fractionation (Leavitt, Flynn, *et al*., 2016), and so the decrease in this system may have played some role in the shifts in the lipid-H compositions we observed (Figure 3). The direction of catabolic S isotope fractionation, lipid-H isotope fractionation, protein and NADH observations are coherent when viewed through the lens of the Rex regulon response to NADH availability at high electron donor flux.

Under electron-donor limitation energy proteins that *significantly increased* in abundance include subunits of both membrane and soluble proteins (Figure 4A). These include the membrane Ech hydrogenase and Rnf complex and an alcohol dehydrogenase (Ald). The Hdr/Flx complex interacts with the ferredoxin reservoir, as well as NAD(H) and alcohol dehydrogenase, which may indicate ferredoxin becomes less reduced under low electron donor availability (Ramos *et al*., 2015), enabling lower flux of reductant to the sulfate reduction machinery, enabling a larger S-isotope fractionation (Figure 1), while simultaneously influencing lipid H-isotope fractionation via the Nfn and NAD(P)(H) reservoirs (Figures 3, 4 and S3).

### 4.2 Changes to ^34^εSO4/H2S and ^2^εlipid/water reflect shifts in catabolism and anabolism, respectively

The cascade of catabolic and anabolic processes *Dv*H operates modulated S and H isotope fractionation profiles, respectively. As predicted, ^34^εSO4/H2S correlated inversely and non-linearly with cell specific sulfate reduction rate, and both respond directly to energy limitation. As csSRR rates slow in response to decreased electron donor flux, the intracellular abundances of key MSR intermediates sulfate, APS and sulfite decrease, as does the isotopic offset between intra– and extracellular sulfate decreases (Figure 2), even as the net fractionation between pooled product sulfide and residual reactant sulfate increases. In a recent study performed with batch-cultures that ran at effectively high csSRR, intracellular sulfate was more than 50 ‰ enriched relative to the extracellular sulfate, even when net fractionation between sulfate and sulfide was an order of magnitude less at 4 ‰ (Sim *et al*., 2017). Together, the prior batch data from Sim and colleagues (2017) and similar observations from these chemostats indicate that the predicted equilibrium isotope effect between sulfate and sulfide is approached intracellularly when sulfate is available in excess of electron donor. The near-equilibrium isotope effect is minimally expressed when csSRR are rapid due to cellular-scale isotope distillation; conversely, the net sulfate/sulfide fractionation values are nearer to equilibrium when csSRR rates and mass-transfer are slow, and less influenced by localized (cellular) distillation. Put differently, the large fractionation predicted from equilibrium theory between sulfate and sulfide is masked by rapid forward reaction rates at the cellular and even enzyme level (Thode *et al*., 1961; Leavitt *et al*., 2014; Sim *et al*., 2023). Large fractionations can only be expressed when net reaction rate slows, and the intracellular isotope effects are summed into the expressed net fractionation values, as has been proposed in some metabolic isotope models (Rees, 1973; Brunner and Bernasconi, 2005; Farquhar *et al*., 2007; Johnston *et al*., 2007; Bradley *et al*., 2011; Sim *et al*., 2019, 2023), but had not been seen until the work by Sim et al. (2017) and now here. Furthermore, because MSR likely operates as a branched metabolic network (Badzjong and Thauer, 1980; Bradley *et al*., 2011), the observed nonlinear relationship between csSRR and ^34^εSO4/H2S (Sim, Bosak, *et al*., 2011; Leavitt *et al*., 2013) remains a challenge to reproduce in models without significant parameterization. Empirical calibrations across a range of organisms and environmental conditions is likely the best option in providing constraints on more nuanced metabolic isotope models.

Lipid-H isotope compositions and electron transfer intermediates also scaled in abundance with electron donor flux and csSRR. The ^2^ε*lipid/water* weighted average in the fastest *Dv*H (13hr) populations were –175 ‰ relative to the slowest (77hr) populations –158 ‰ (Figure 3B), with lipid-specific isotope pattern (i.e., chain length and iso-/anteiso-/*n*-) followed prior observations for fatty acids from sulfate reducing bacteria (Campbell *et al*., 2009, 2017; Dawson *et al*., 2015; Leavitt, Flynn, *et al*., 2016; Leavitt, Venceslau, *et al*., 2016; Osburn *et al*., 2016; Leavitt *et al*., 2017, 2019). In addition to fatty acids, more membrane-bound electron transfer intermediates menaquinone-5 and –6 are present in 13hr relative to 77hr cells (Figure S2B). Beyond *Dv*H, the trend of more depleted lipids at higher growth rates is consistent with batch experiments utilizing other sulfate reducers across a range of carbon and energy sources (Leavitt, Flynn, *et al*., 2016; Osburn *et al*., 2016). We also provide coarse estimates of the soluble NAD(P)(H) reservoirs, which were all higher in the faster cells (Figure S2A), where we also observed increased NfnB expression at higher energy flux (conversely less NfnB at lower energy flux, Figure 4B). This is broadly consistent with two studies in a sulfate reducer and fermenter, where genes (*nfnAB*) encoding the NADP(H)/NAD(H) electron-bifurcating transhydrogenase, were genetically disrupted, which forced the organisms to grow more slowly than the respective wildtypes, and led to less ^2^εlipid/water fractionation (Leavitt, Flynn, *et al*., 2016; Leavitt *et al*., 2017). These results are consistent within anaerobic bacteria that utilize the Nfn transhydrogenase, where the broader observation of less lipid/water H-isotope fractionation at lower growth rates, perhaps due to lower energy flux (Leavitt, Flynn, *et al*., 2016; Osburn *et al*., 2016; Leavitt *et al*., 2017). These observations across experiment type indicate a central role for energy availability in setting the net ^2^εlipid/water values, primarily via the recording of intracellular NADPH/NADH (im)balances in lipid-H isotope compositions (Wijker *et al*., 2019). Future works might track Domain or metabolism specific lipids and lipid-H isotope values across environmental energy gradients, particularly in systems where direct cultivation or activity assays are not practical or yet possible, e.g. deep subsurface sediments, subglacial lakes, and deep fracture fluids (Lever *et al*., 2015; Drake *et al*., 2018; Bradley *et al*., 2020).

### 4.3 Framework to interpret the response of sulfate reducers to energy limitation

A theme emerges that ties together *D. vulgaris*’ response to different reductant availability as they are expressed in inorganic S and organic H isotopes, protein, and metabolites.

***High energy flux*** – When electron and carbon source flux is highest (13hr population), DvH cells have to transfer reducing equivalents onto sulfate with maximum efficiency. In order to process incoming reducing equivalents, cells in the 13hr chemostats maintain high stocks of the MSR intermediates APS and sulfite (Figure 2B), while also accumulating a larger intracellular pool of the terminal electron acceptor sulfate (Figure 2B). This is also seen in the higher abundance of AprAB and QmoAB proteins in the 13hr cells (Figure 3B), and increased menaquinone levels in the faster growing cells (Figure S2B). This enables efficient electron transport to and through the membrane and the rapid reduction of sulfate, which then limits the expressed S isotope fractionation between sulfate and sulfide (Figures 1 and 2A). The larger standing pools of electron-transfer (MK) and reducing equivalents in the faster 13hr cells is broadly consistent with a higher availability of electrons under the higher reductant flux regime. This indicates that the DvH populations that experienced larger electron donor fluxes were able to build transmission capacity, which again enabled faster sulfate respiration. This is consistent with the protein and S-metabolite data, where higher abundances of the QmoAB electron transfer proteins, more intracellular sulfate sulfite and APS at the faster csSRR, all converge to generate smaller net S isotope fractionation values and larger net H isotope values.

***Low energy flux* –** During severe electron donor-limitation, APS reductase and QmoAB were less expressed and likely csSRR decreased, which are reflected in the increased ^34^εsulfate/sulfide. The control energy flux exerts on csSRR and ^34^εsulfate/sulfide appears to be mediated by APS reductase abundance. While it is possible csSRR could be under post-translational control, the near constant ratios of APS:sulfite in both the 13hr and 77hr conditions suggests that increased flux through that step of sulfate reduction is indeed controlled by APS reductase abundance. Still, Further testing of this hypothesis will require independent genetic control of AprAB abundance.

Broadly, these findings agree with two recent batch-culture studies by Sim and colleagues (Sim *et al*., 2017, 2019). With energy least available in the 77hr chemostats, electron donor overburden was not as much of a challenge, intracellular sulfate concentrations and sulfate reduction metabolites were lower (Figure 2B), and the S-isotope offset between extra– and intracellular sulfate was eliminated. Enzymes needed to reduce sulfate to sulfite were in lower demand (cf. AprAB, QmoAB), whereas proteins that enable more efficient energy acquisition from the carbon and electron donor (Ldh, Por, Pta, Ack) were then in higher demand, as were those needed to couple electron transfer to membrane redox potential (cf. Ech, Tmc, Rnf) and to facilitate cytoplasmic electron transfer (Flx/Hdr, certain Adh’s) (Figure 5). These observations are consistent with the notion that as biochemical reducing equivalents became less available due to lesser lactate influx, the DvH cells increased the abundance of the key, perhaps high affinity, Ech hydrogenase to maintain proton motive force in the face of lower energy flux. The large increase in the membrane energy metabolism Rnf complex (Figure 4A), which links the NADH and ferredoxin pools, is likely a reflection of the need for energy to continue through the system at lower flow rates. Consistent with this concept, we observe more membrane bound and soluble electron transfer machinery at the lowest power regime (77hr; specifically, Ech, Rnf, Hdr/Flx, Por, Aor), as compared to more sulfate-activation machinery in the intermediate (26 hr), and especially fastest (13hr) conditions –-specifically, more ApsR and Qmo (Figures 4 and 5). Taken together, it is most likely that that APS reductase expression and the Rex regulon most modulate variation in ^34^εSO4/H2S and ^2^εlipid/water across the different energy limited conditions in these experiments

**Figure 5.**
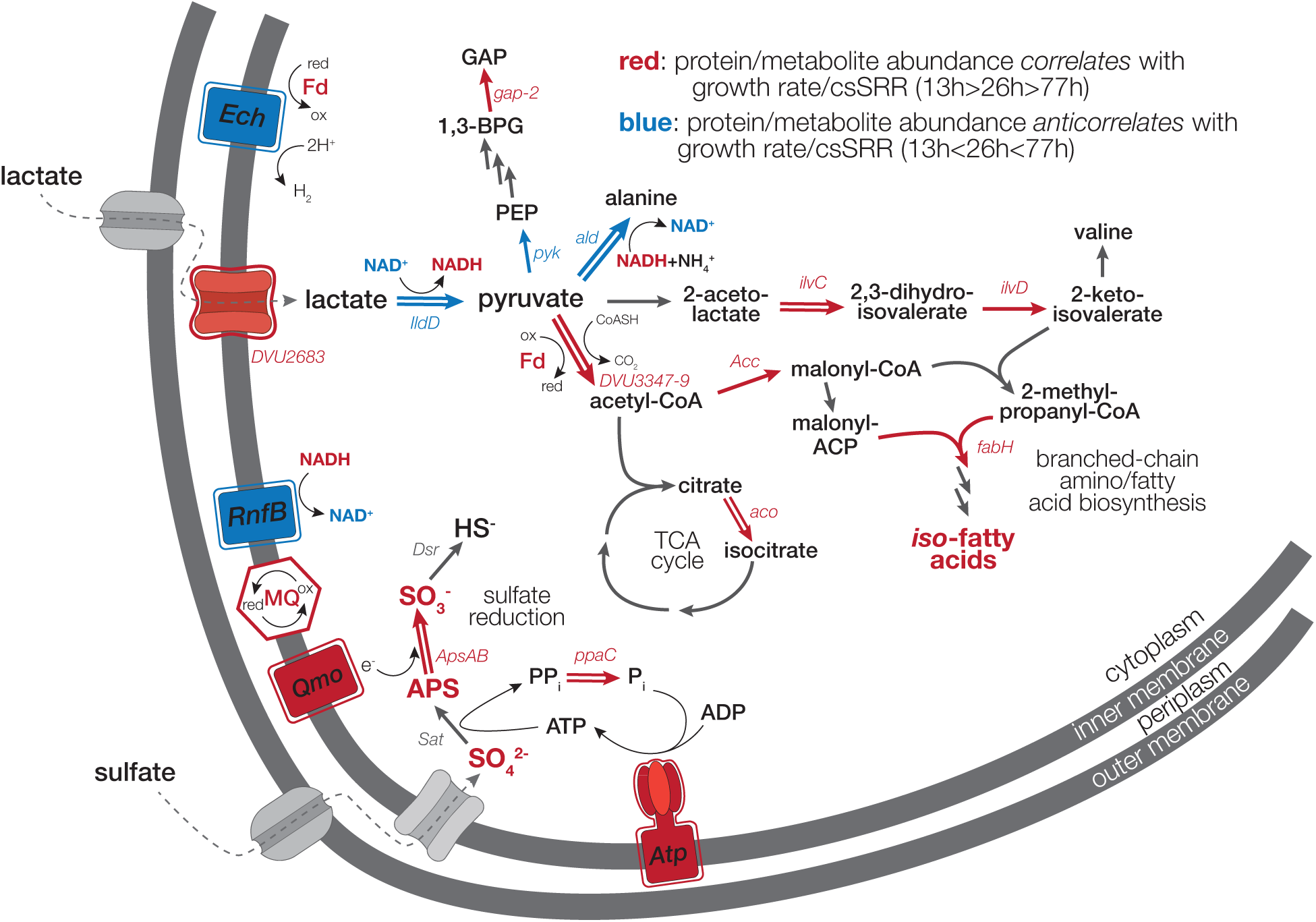
A schematic summary of observed protein and metabolite abundance changes in *Dv*H with rate modulation in lactate-limited chemostats. This summary focuses on sulfur, carbon and energy metabolism. The abundance of proteins/metabolites highlighted in red correlates with csSRR (and inversely correlated with ^34^ε_SO4/H2S_); the abundance of blue anti correlated with csSRR (and correlated with ^34^ε_SO4/H2S_). For proteins, a double arrow or outline indicates that the abundance difference between the 13 hr and 77 hr rates is significant at q <0.05.

***Remaining Challenges*** – While the approaches here shed much light on the relationship between thermodynamic drive and net catabolic isotope fractionations, it remains unclear if the major intracellular electron carriers change in response to energy flux. It was predicted that different electron carriers are utilized under different power regimes (Wing and Halevy, 2014; Wenk *et al*., 2017), which may include: ferredoxin (–398 mV); two forms of flavodoxin (–371 and –115 mV); cytochrome *c*3 (–290 mV); menaquinones (–74 mV) and rubredoxin (–57 mV) – midpoint potentials from Thauer (Thauer *et al*., 1977)*th*, but this was not confirmed here. Wenk and colleagues (2017) focus on rubrerythrin (+23mV), though to-date this protein is known only to play a role in oxygen detoxification (Coulter *et al*., 1999; Kurtz Jr, 2006). We do observe small shifts in the abundance of the menaquinones themselves, and in the cellular NAD(P)(H) pools (Figure S2A). Indeed, the proteomic approach here does not resolve such changes, which may occur at the scale of electron flux to/through key redox carriers (e.g. ferredoxin, menaquinones, and NAD(P)H)). In such a scenario, the total number of and average redox state of electron transfer intermediates sets the capacity for energy transduction, which the cell increases or decreases as necessitated by environmental conditions.

Though shifts in proteomic and metabolite profiles indicate changes in the flow of energy through the electron transport chain, the extent to which shifts in electron carriers per prior predictions does occur (Wenk *et al*., 2017; Sim *et al*., 2019; Bertran *et al*., 2020), particularly at extremely low energy fluxes, remains experimentally under-constrained. Follow up studies should work to directly quantify the total abundance and the oxidized/reduced ratios of the key players under a range of energy (and ultimately power) limitations. To further explore the relationship between net S-isotope fractionation, intracellular isotopic compositions, proteins, and metabolic flux-balance, we suggest the chemostat and associated analytical approaches employed herein be repeated at even more extreme energy (or sulfate, or nutrient) limitations as have been tested previously (Sim, Bosak, *et al*., 2011; Leavitt *et al*., 2013).

## 5. CONCLUSIONS

The coordinated shifts in proteins and metabolites underpins the non-linear relationship between the observed csSRR and ^34^εSO4/H2S (c.f. Figures 1, S4). We show that DvH modulates the composition of its membrane, electron carriers, and redox active proteins in response to the different energy flux regimes imposed upon it (Figures 2, S2, 3 and 4). Collectively, this modulates the expressed S and H isotope fractionation values for catabolism and anabolism, respectively, each as they relate to central energy metabolism and csSRR, presenting unique ‘isotopic phenotypes’ (Pellerin *et al*., 2015, 2018). The availability of energy in any environment dictates the ecological and geochemical responses of the local microbial community. In chemostat experiments with the well-studied sulfate reducer *Desulfovibrio vulgaris* Hildenborough, the lipid, protein and redox active metabolites respond coherently to differences in power availability, which yield distinct and predictable shifts in cell-specific respiration rate, lipid/water H and sulfate/sulfide S isotope values. While it remains to be understood how precisely the biochemical reactions sum and translate into the inverse relationship between respiration rate and specifically S-isotope fractionation, it is clear that the cells respond in real time to the influx of energy and can modulate a wide range of their electron transport chain and shift the standing stocks of respiration intermediates.

Looking forward, this work will inform our understanding of power availability, such that cells operate as capacitors to maintain net forward reaction progress in the core of sulfate reduction, while tuning intracellular energy demands in proportion to environmental conditions. To incorporate these findings into metabolic isotope model, the quantities of both MSR reaction intermediates and electron transfer proteins must be carefully calibrated in response to a much wider range of power availabilities reflecting those in natural systems, where life operates close to the thermodynamic limit (Hoehler and Jørgensen, 2013; Hoehler *et al*., 2023), tuned to incorporate natural abundance stable isotopic constraints, as well as with a more diverse array of sulfate reducers or metabolisms operating near the thermodynamic limit. The combination of experimental and analytical approaches employed here offers a fresh look at the cellular mechanisms that underpin Earth’s S and C cycles through the lens of dissimilatory sulfate reduction. The merger of careful physiological work with isotopic analyses will enable us to calibrate metabolic isotope models, both in the MSR system, as well as microbial metabolisms that regulate elemental cycles at a planetary scale.

## 6. DATA AVAILABILITY

**Proteomic data**: Proteomic mass spectral data are available via proteomeXchange under accession PXD027511 and the MassIVE repository (massive.ucsd.edu) under accession MSV000087869 [*username for reviewers*: MSV000087869_reviewer / password: desulfo34S].

**Other Data**: All experimental and analytical data are permanently available in the supplemental Dataframes, that will be permanently available on FigShare [Private Link for Reviw: https://figshare.com/s/249464bd6a761d5c7889].

## Acknowledgements

We thank S. Moore for assistance with bulk sulfur isotope analyses (Fike lab, WashU), M. Seuss for assistance with lipid-H isotope analyses (Bradley lab, WashU); X. Feng (Dartmouth) and M. Osburn (Northwestern) for water H-isotope analyses; A. Sessions and J. Adkins (CalTech) for access to HPLC-ICP-MS. Metabolite analyses were performed by the Proteomics & Mass Spectrometry Facility at the Danforth Plant Science Center (St. Louis, MO, USA). Funding was provided: by NASA Exobiology Award 13-EXO13-0082 (ASB, DF, WDL, JW), NSF-EAR Award 1928309 (WDL), Washington University in St. Louis Department of Earth & Planetary Sciences Fossett Fellowship (WDL), the Walter and Constance Burke Fund at Dartmouth College (WDL), and the Fulbright – Bunge & Born – Williams Foundation Scholarship Program (FJB), Fundação para a Ciência e Tecnologia (Portugal) through R&D unit MOSTMICRO-ITQB (UIDB/04612/2020 and UIDP/04612/2020) and LS4FUTURE Associated Laboratory (LA/P/0087/2020), NSF GRFP [2017250547] (SP).

## MAIN FIGURES & TABLE

Figure 1. Rate vs. net Fractionation;

Figure 2: Sulfur reservoirs and isotopes vs. rate

Figure 3. Lipids and H-isotopes.

Figure 4. Energy proteins vs. Rate.

Figure 5. Metabolism Cartoon summary

Table 1. Differentially expressed proteins with rate, by COG

## SUPPLEMENTAL FIGURES

Figure S1. Chemostat OD’s vs. time.

Figure S2: Soluble and membrane associated electron and hydride carrier abundances.

Figure S3. Proteomics Summary.

Figure S4: S-isotope & energy protein shift co-plotted. & Energy Proteomic shifts summary re-plotted at scale.

## SUPPLEMENTAL DATA AVAILABILITY

**Other Data**: All experimental and analytical data are permanently available in the supplemental Dataframes, that will be permanently available on FigShare [Private Link for Review: https://figshare.com/s/249464bd6a761d5c7889].

**Dataframe 1: Chemostat data; S isotopes & metabolites; cellular metabolites; cell counts.**

**Dataframe 2. Lipid H-isotopes.**

**Dataframe 3. Proteomics.**

**Figure S1.**
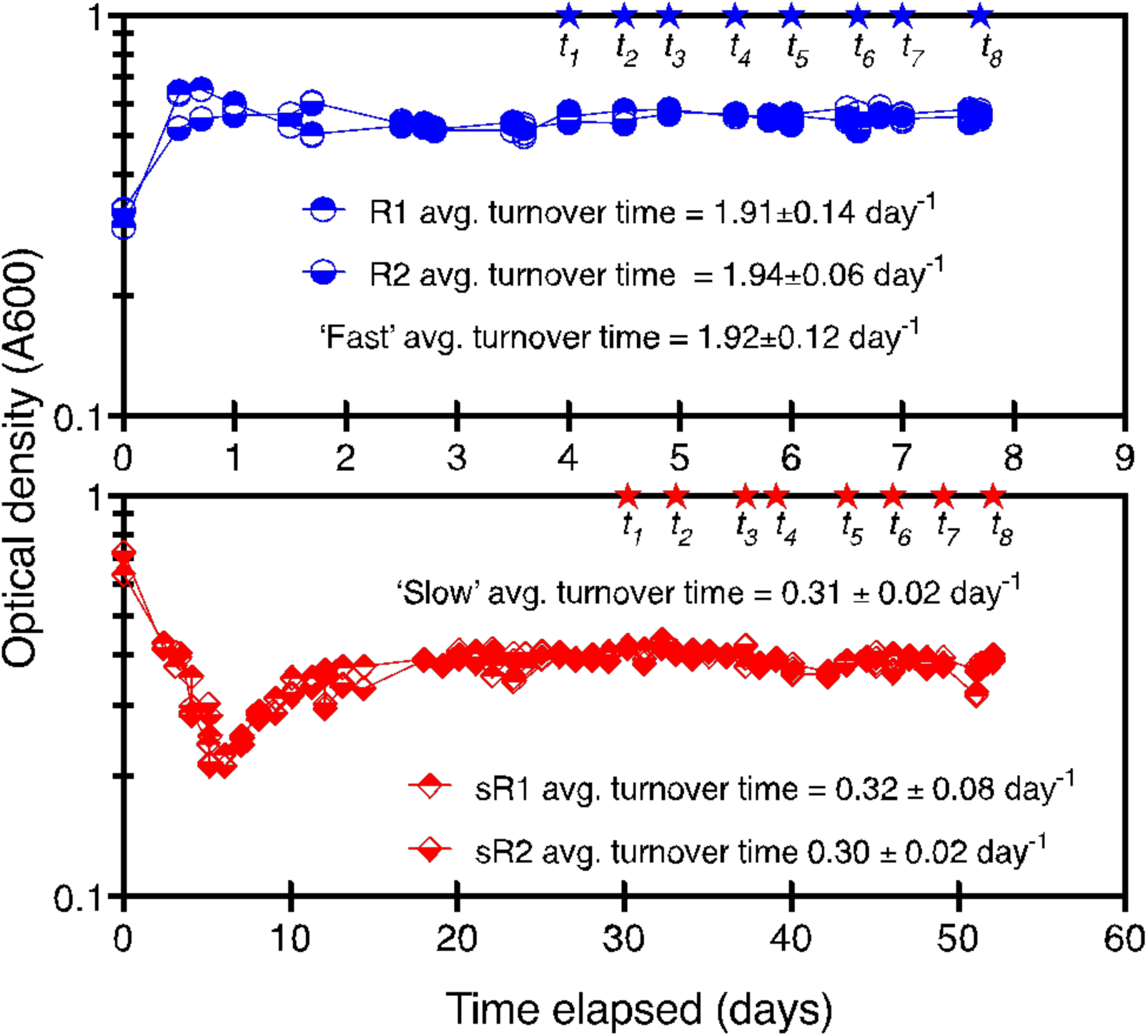
Visual display of steady-state experiments. Optical density of each bioreactor over the course of each chemostat experiment (circle and diamonds), with the eight sampled time points (stars). The top panel is from the fast reactors (∼13 hr doubling time, proportional to 1.91 to 1.94 per day), whereas the bottom panel are from the slow turnover time reactors (∼77 hr doubling time, proportional to 0.30 to 0.32 per day). The intermediate rate of ∼26hr data is from Leavitt et al. 2019.

**Supplementary Figure S2.**
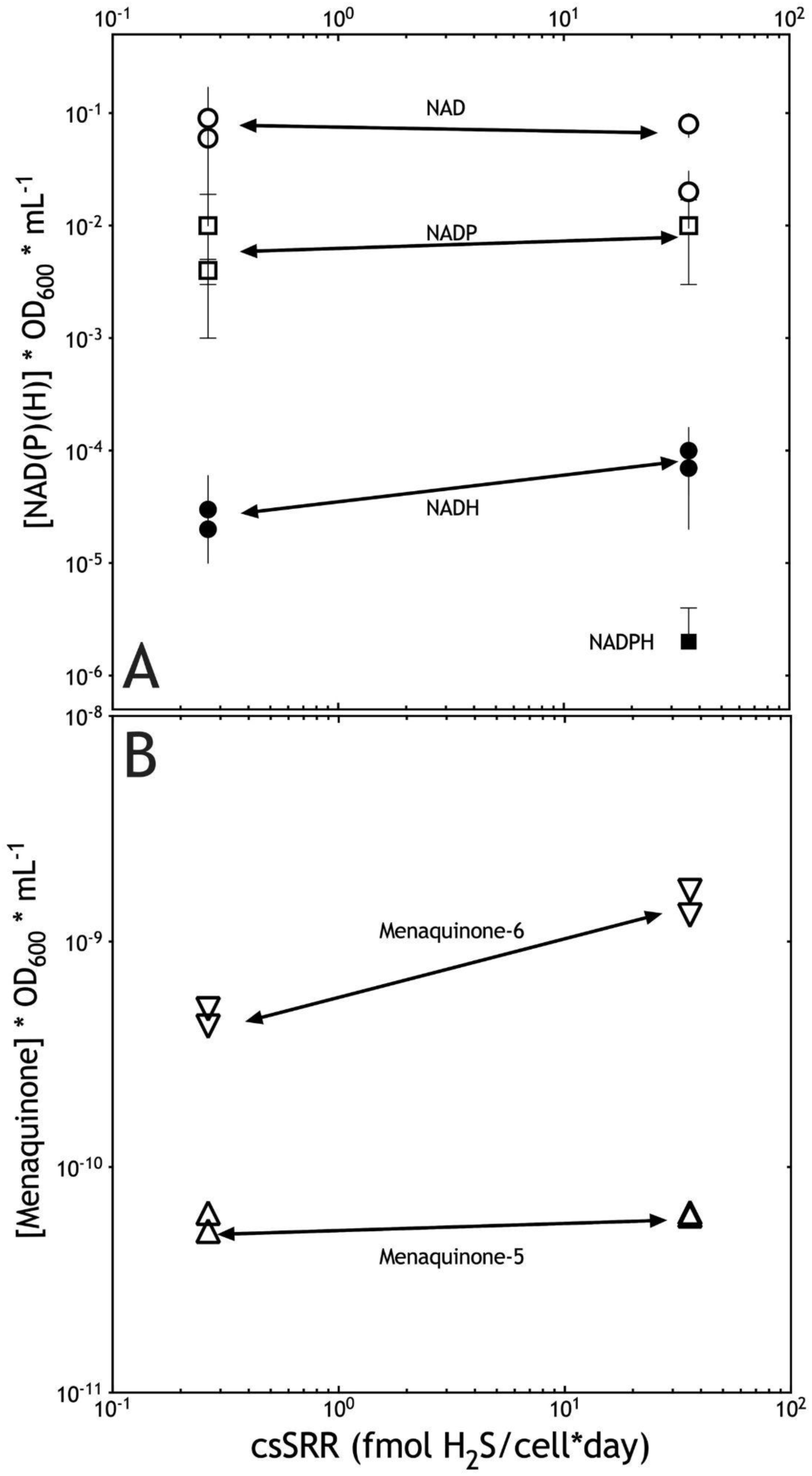
Cellular electron transfer intermediates from the slowest (77hr) and fastest (13hr) chemostat grown *Dv*H populations plotted versus csSRR. (A) soluble intracellular NAD(P)(H) hydride carriers, where each pool is normalized to a unit of biomass (OD600). (B) Membrane associated menaquinone’s (MK’s) 5 and 6, also normalized to biomass and measurements standards MK-4. These measurements were not collected in the Leavitt et al. (2019) study, and so are not available for 26hr *Dv*H chemostat population.

**Supplemental Figure S3.**
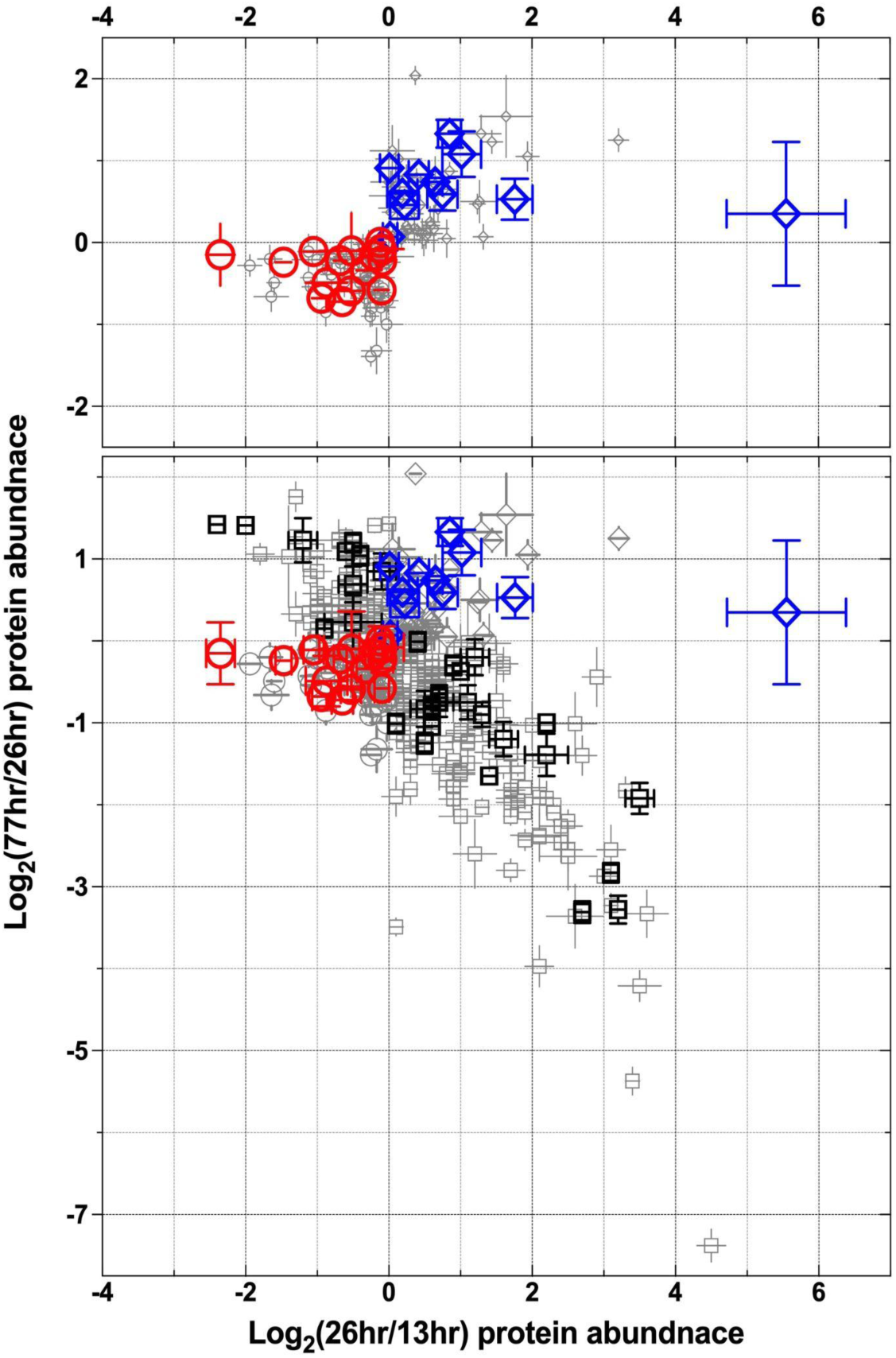
Proteomics Summary. COG-identified *energy proteins* from *Dv*H whose abundance scales positively with csSRR (red circles, most in 13hr) or inverse with csSRR (blue diamonds, most in 77hr), or outliers that are most or least abundant in the intermediate rate (black squares, most or least in 26hr). All other proteins whose abundance scaled significantly and positively with rate (gray circles), inverse to rate (gray diamonds), or most/lease in the intermediate rate (gray squares). The top panel only shows significantly different energy proteins (blue and red symbols are those featured in Figure 4 and S4), the bottom panel shows all proteins detected at all rates that showed statistically significant patterns. Symbols are the same across panels.

**Figure S4.**
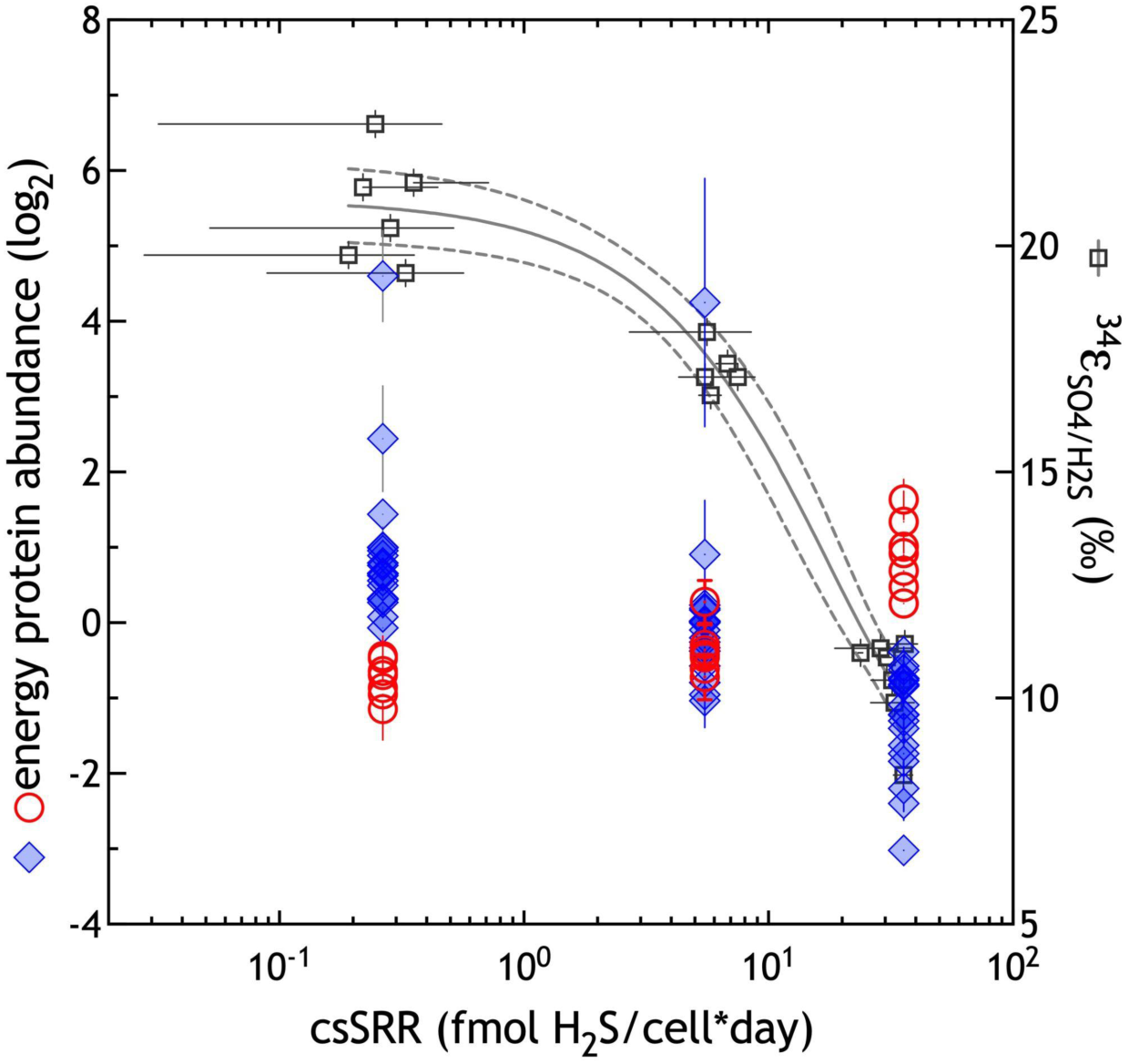
Chemostat S isotope fractionation and energy proteomics summary. These are the same data as in Figure 1B (black squares, right y-axis), and Figure 4A (blue diamonds) and 4B (red circles), here plotted with the same (left) y-axis scale.

